# The first draft genomes of the ant *Formica exsecta*, and its *Wolbachia* endosymbiont reveal extensive gene transfer from endosymbiont to host

**DOI:** 10.1101/436691

**Authors:** Kishor Dhaygude, Abhilash Nair, Helena Johansson, Yannick Wurm, Liselotte Sundström

## Abstract

The wood ant *Formica exsecta* (Formicidae; Hymenoptera), is a common ant species throughout the Palearctic region. The species is a well established model for studies of ecological characteristics and evolutionary conflict. In this study, we sequenced and assembled draft genomes for *Formica exsecta* and its endosymbiont *Wolbachia*. The draft *F. exsecta* genome is 277.7 Mb long; we identify 13,767 protein coding genes for which we provide gene ontology, and protein domain annotations. This is also the first report of a *Wolbachia* genome from ants, and provides insights into the phylogenetic position of this endosymbiont. We also identified multiple horizontal gene transfer events (HGTs) from *Wolbachia* to *F. exsecta*. Some of these HGTs have also occurred in parallel in multiple other insect genomes, highlighting the extent of HGTs in eukaryotes. We expect that the *F. exsecta* genome will be valuable resource in further exploration of the molecular basis of the evolution of social organization.

## Introduction

Adapting to changes in the environment is the foundation of species survival, and is usually thought to be a gradual process. Genomic changes, such as single nucleotide substitutions play key roles in adaptive evolution, although few mutations are beneficial. Besides nucleotide substitutions, other structural and regulatory units, such as transposable elements (TEs) and epigenetic modifications, can also act as drivers in adaptation (González et al., 2010; Rostant, Wedell & Hosken, 2012; Casacuberta & González, 2013). Genetic material can also be acquired from other organisms by means of horizontal gene transfer (HGTs), and this can also lead to novel adaptive traits (Schönknecht, Weber & Lercher, 2014; Wybouw et al., 2016). Both mutations and HGTs can drive rapid genome evolution (Dunning Hotopp, 2011; Boto, 2014). Horizontal gene transfers have been reported in many taxa, most commonly from bacteria to eukaryotes (Dunning Hotopp, 2011), plants (Yue et al., 2012; Matveeva & Lutova, 2014), fungi (Rolland et al., 2009; Fitzpatrick, 2012; Bruto et al., 2014), but the underlying mechanisms that underpin horizontal gene transfer events, and mode by which bacterial genetic material is integrated into the eukaryote genome are not well understood.

Many cases of horizontal gene transfer from bacteria to eukaryotes involve intracellular endosymbionts, which are maternally transmitted through oocytes (Werren, 1997]; Ferree et al., 2005). The most common examples of endosymbiont to host horizontal gene transfers involve the bacterium *Wolbachia*, a well described intracellular, maternally inherited gram-negative bacterium known to infect over 40% of the investigated insect species (Werren, 1997; Werren, Baldo & Clark, 2008). *Wolbachia* infection is also prevalent in filarial nematodes, crustaceans, and arachnids (Cordaux, Michel-Salzat & Bouchon, 2001; Fenn et al., 2006; Goodacre et al., 2006). *Wolbachia* host interactions can be mutualistic or pathogenic (Moya et al., 2008). A number of ecdysozoan genomes have been reported to contain chromosomal insertions originating from *Wolbachia*, including the mosquito *Aedes aegypti* (Klasson et al., 2009a; Woolfit et al., 2009), the longhorn beetle *Monochamus alternatus* (Aikawa et al., 2009), filarial nematodes of the genera *Onchocerca, Brugia*, and *Dirofilaria* (Fenn et al., 2006; Hotopp et al., 2007), parasitoid wasps of the genus *Nasonia*, the fruit fly *Drosophila ananassae*, the pea aphid *Acythosiphon pisum* (Nikoh & Nakabachi, 2009; Nikoh et al., 2010), and the bean beetle *Callosobruchus chinensis* (Kondo et al., 2002). Although most of the transferred DNA is probably nonfunctional in the host genome (Kondo et al., 2002; Hotopp et al., 2007; Nikoh et al., 2008), some of the transferred genes are functional (Klasson et al., 2009a). These genes are expressed in specific tissues, are subject to purifying selection, and are involved in processes such as protein synthesis inhibition, membrane transport and metabolism (Hotopp et al., 2007; Woolfit et al., 2009; McNulty et al., 2013).

Infection with *Wolbachia* is widespread in Hymenoptera. Most hymenopteran *Wolbachia* infections have the cytoplasmic incompatibility phenotype (Werren & Windsor, 2000), which leads to reproductive incompatibility between infected sperm and uninfected eggs. Wenseleers et al. (1998) showed that 25 out of 50 species of ants in Java and Sumatra screened positive for one strain of *Wolbachia*. By contrast, a study on a single Swiss population of the ant *Formica exsecta*, found that all the ants tested were infected with four or five different strains of *Wolbachia* (Keller et al., 2001; Reuter & Keller, 2003).

The aims of this study are to test whether horizontally transferred genetic elements exist in the genome of the ant *Formica exsecta*, and to describe the genomic organization of any such elements. The genus *Formica* is listed by the Global Ant Genome Alliance (GAGA) as one of the high-priority ant taxons to be sequenced (Boomsma et al., 2017; http://antgenomics.dk/), owing to its key taxonomic position, and the ecological and behavioral data that are available for the species. To date, no genome sequence is available for this genus.

Our study population of *F. exsecta*, located on the Hanko peninsula, Southwestern Finland, has been monitored since 1994, and data on demography, genetic structure, and ecology are available (Sundström, Chapuisat & Keller, 1996; Sundström, Keller & Chapuisat, 2003; Haag-Liautard et al., 2009; Vitikainen, Haag-Liautard & Sundström, 2015). Based on genetic data on colony kin structure most (97%) of the approximately 200 colonies are known to have a single reproductive queen, mated to one or more (usually two) males (Sundström, Chapuisat & Keller, 1996; Sundström, Keller & Chapuisat, 2003; Haag-Liautard et al., 2009; Vitikainen, Haag-Liautard & Sundström, 2015). We report the whole genome sequencing of this species, and the draft genome sequence of its associated cytoplasmic *Wolbachia* endosymbiont (wFex). We further report the presence of multiple extensive insertions of *Wolbachia* genetic material in the host genome, and compare the HGTs insertions discovered in the assembled draft genome to other genomes, to understand the pattern of HGT events between endosymbiont and host. We analyze in detail the genomic features of *F. exsecta* along with its endosymbiont *Wolbachia*, and discuss our findings in the light of genome evolution in *Wolbachia* and its host.

## Materials and Methods

### Sample collection and genome sequencing

We selected one single-queen colony from our study population on the island Furuskär (F162), and collected 200 adult males from this colony. We used males because in Hymenoptera these arise through arrhenotoky (Normark, 2003) and are haploid (Crozier, 1975), meaning that a pool of males together are representative of the diploid genome of their mother. DNA extraction was done from testis, which contains sperm cells and organ tissue, to avoid contamination by gut microbiota. We used a Qiagen Genomic-tip 20/G extraction kit according to the manufacturer’s protocol. For Illumina sequencing we constructed three small insert paired-end libraries (insert sizes of 200 bp, 500 bp, 800 bp), and four mate pair (large insert paired-end) libraries (insert sizes of 2 kb, 5 kb, 10 kb and 20 kb), each containing DNA from 15-50 pooled males. Libraries were prepared using protocols recommended by the manufacturers. Sequencing was done at the Beijing Genomics Institute (BGI) using HiSeq2000, which produced a total of 99.97 GB of raw data (Table 1).

**Table 1:**
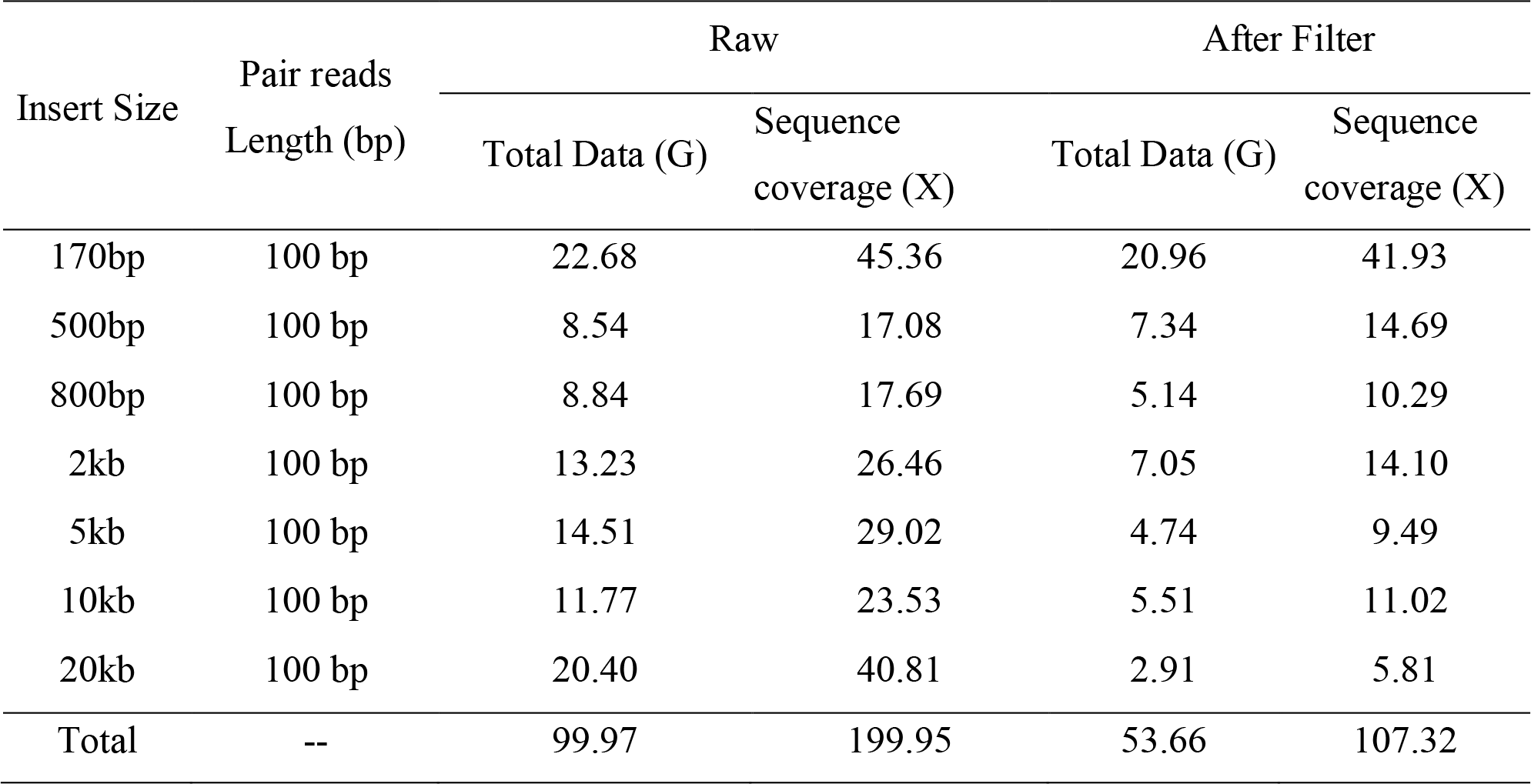
Summary statistics for the raw sequencing data, before and after filtering reads. “Coverage depth” was calculated based on the estimated assembled genome size (300 Mb).

### Genome assembly

We assembled the *F. exsecta* genome using SOAPdenovo2 version 2.04 (Xie et al., 2014) in three main steps. First, a de Bruijn graph was constructed using short length insert library reads with default parameters (k-mer value of 45), to construct the contigs. The initial contig assembly contained 104,190 contigs with an N50 size of 22,328 bp, and total length of 276.23 Mb of sequence, at an average depth of coverage of 47.37X. Second, all individual reads were realigned onto the contigs. Because reads are paired, they can aid with scaffolding: The number of reads supporting the adjacency of each pair of contigs was calculated and weighted by the ratio between consistent and conflicting paired ends. Scaffolds were constructed in a stepwise manner using libraries of increasing sizes from 500bp insert size paired-end reads up to mate-pair of 5 kb insert size. 80,473 contigs could not be placed in scaffolds. These are highly similar repetitive sequences, since the cd-hit-est tool (Huang et al., 2010) showed that 43% of these contigs clustered together at 80% of the sequence length. Third, sequencing gaps in the scaffolds were closed with the two mate-pair libraries (Insert size 10 kb and 20 kb). Overall, these steps produced an initial assembly with an N50 scaffold length of 949,634 bp, and a total length of 289,843,734 bp with each scaffold longer than 200 bp.

We used blobology v1.0 (Kumar et al., 2013) to generate taxon-annotated GC coverage (TAGC) plots of scaffolds in the genome assembly, which can help to identify bacterial contamination (Supplementary Figure S1). The scaffolds for the TAGC plot were successfully annotated to the taxonomic order based on the best blast match to the NCBI nt database (O’Leary et al., 2016). This analysis revealed that 74 scaffolds matched the endosymbiotic bacterium *Wolbachia.* Sixty-nine of these scaffolds were removed as we concluded that they are part of the *Wolbachia* genome (see analysis below), but five contigs were retained in the final assembly for *F. exsecta* as they contained both *Wolbachia* and ant sequences. Following this curation, the final draft genome assembly was 277.7 Mb long with an N50 value of 997,654 bp and 36% Guanine-cytosine (GC) content (Table 2).

**Table 2:** Genome assembly statistics for *F. exsecta* and its *Wolbachia* endosymbiont.

| **Genome Assembly** | ***Formica exsecta*** | **FE *Wolbachia*** |
| --- | --- | --- |
| **Stats** | **Genome** | **endosymbiont Genome** |
| Total length | 277719392 (277 MB) | 3096460 (3.09 MB) |
| Total contigs | 14617 | 69 |
| Contigs (>= 1000 bp) | 3136 (98.24% genome) | 68(99.97% genome) |
| Contigs (>= 50000 bp) | 545 (89.59% genome) | 22(75.48% genome) |
| N50: | 997654 bp | 104167 bp |
| N75: | 318356 bp | 54296 bp |
| L50: | 73 | 11 |
| L75: | 185 | 22 |
| GC (%) | 36.00 | 35.13 |

### Genome assembly of *Wolbachia*

All 25 published *Wolbachia* genomes were obtained from the NCBI database (O’Leary et al., 2016). We aligned the 74 scaffolds from the initial *F. exsecta* assembly that matched with *Wolbachia* against these genomes using MUMmer 3.23 (Kurtz et al., 2004), and inspected the alignments manually. Sixty-nine of the 74 scaffolds matched completely to *Wolbachia* genomic regions. These 69 scaffolds represented 3.09 Mb total, with a N50 value of 104,167 bp, henceforth we refer to this group of scaffolds as “the *Wolbachia* endosymbiont genome of *F. exsecta*” (wFex).

The remaining five scaffolds each contained several interspersed fragments with similarity to *Wolbachia* genomes, whereas other parts of these scaffolds had high similarity to genomes of ants. Furthermore, the sequencing coverage of these scaffolds was similar to the *F. exsecta* scaffolds, rather than to the *Wolbachia* scaffolds. Finally, detailed inspection of these scaffolds in a genome browser showed no change in sequencing depth where we identify the interspersed fragments with similarity to *Wolbachia*, which would be expected for erroneous chimeric assembly (Lasken & Stockwell, 2007). These data thus suggest that fragments of *Wolbachia* were horizontally transferred to the *F. exsecta* genome. To corroborate these results with independent approaches, we re-assembled the raw sequencing data with two additional independent algorithms that we expect would make different types of assembly errors than SOAPdenovo. The first software, Velvet version 1.2.09 (Zerbino & Birney, 2008), is also based on a de Bruijn graph; the second, SGA version 0.10.5 (Simpson & Durbin, 2012) is based on a string graph. Both resulting assemblies confirmed the patterns we had seen, and validate the idea that the five SOAPdenovo scaffolds containing sequence with similarity to both ants, and *Wolbachia* represent horizontal gene transfers from *Wolbachia* to *F. exsecta*.

We further compared the sequences of the horizontally transferred fragments in the five SOAPdenovo scaffolds against the NCBI (nr/nt) database (O’Leary et al., 2016), using blast 2.2.27 (Altschul et al., 1990) to determine whether these fragments may have also undergone horizontal gene transfer in other arthropod genomes. We performed analogous searches on ant genomes present in the NCBI, and the Fourmidable databases (Wurm et al., 2009). When a positive match with any other ant or arthropod genomes was found, the exact location of the insertion was determined, and compared with that of *F. exsecta.* Finally, the five scaffolds were also compared to the *F. exsecta* transcriptome (Dhaygude et al., 2017), using blastn 2.2.27, to assess similarity with expressed sequences.

### Quantitative assessment of genome assemblies

The quality of the genome assembly is crucial, as it defines the quality of all subsequent analyses that are based on the genome sequences. We explored multiple assembly options (data not shown), and used two methods to assess assembly quality and robustness in order to select the highest quality assembly. First, we evaluated genome contiguity (number and length of contigs) using Quast 3.2 (Gurevich et al., 2013) to assess whether our newly assembled draft genome is comparable to published ant genomes (Favreau et al., 2018) based on assembly statistics (N50,N90). Second, we used core gene content-based quality assessment using CEGMA 2.4 (Parra et al., 2007) to ascertain that the 248 most highly conserved eukaryotic proteins are present in our genome assembly. We also compared genes present in our genome assembly to single-copy orthologs across four lineage-specific sets (Eukaryota (303 genes), Insecta (1,658 genes), Arthropoda (2,675 genes), and Hymenoptera (4,415 genes)) using the BUSCO 1.1(Simão et al., 2015). In addition, we compared the *F. exsecta* genome with 13 other ant genomes, *Camponotus floridanus, Atta cephalotes, Acromyrmex echinatior, Cardiocondyla obscurior, Cerapachys biroi, Lasius niger, Linepithema humile, Monomorium pharaonis, Pogonomyrmex barbatus, Vollenhovia emeryi, Wasmannia auropunctata, Harpegnathos saltator*, and *Solenopsis invicta* (Wurm et al., 2009), using BUSCO. We report BUSCO quality metrices for the *F. exsecta* genome. (Table 3).

**Table 3:** BUSCO quality metrics for the *F. exsecta* genome and the *Wolbachia* endosymbiont of *F. exsecta (wFex)* genome assembly

| BUSCO metric | *F. exsecta* genome |  |  |  | *FE Wolbachia*<br>*endosymbiont Genome (wFex)* |  |
| --- | --- | --- | --- | --- | --- | --- |
|  | Eukaryota | Insecta | Arthropoda | Hymenoptera | Bacteria | Proteobacteria |
| Complete | 299 (98.7%) | 1634(98.5%) | 2549(95.29%) | 4249 (96.2%) | 107 (72.30%) | 158 (71.49%) |
| Complete and single copy | 283 (93.4%) | 1572 (94.8%) | 2446(91.44%) | 4151 (94.0%) | 35 (23.65%) | 55 (24.88) |
| Complete and duplicated | 16 (5.3%) | 62 (3.7%) | 103 (3.86%) | 98 (2.2%) | 72 (48.65%) | 103 (46.60%) |
| Fragmented | 1 (0.3%) | 15 (0.9%) | 195 (7.29%) | 123 (2.8%) | 9 (6.08%) | 11 (4.97%) |
| Missing | 3 (1.0%) | 9 (0.6%) | 34 (1.27%) | 43 (1.0%) | 32 (21.62%) | 52 (23.52%) |
| Total | 303 (100%) | 1658 (100%) | 2675 (100%) | 4415 (100%) | 148 (100%) | 221 (100%) |

The quality of the *Wolbachia* endosymbiont genome was quantified with a similar approach, where we used BUSCO to examine the presence of Universal Single-Copy Orthologs of the Bacteria (148 genes), and the Proteobacteria (221 genes) lineages (Table 3). We also used BUSCO to compare the wFex genome with four other *Wolbachia* genomes, including the *Wolbachia* endosymbionts of *Drosophila simulans (wRi), Culex quinquefasciatus (wPip), Drosophila melanogaster (wMel)*, and *Drosophila simulans (wNo)*.

### Gene prediction

We combined several publicly available data sets and computational gene prediction tools to establish an Official Gene Set (OGS) for the *F. exsecta* genome. First, we used the MAKER version 2.28 pipeline (Cantarel et al., 2008; Holt & Yandell, 2011), to derive consensus gene models from Augustus version 3.1.0 (Stanke & Morgenstern, 2005), SNAP version 2016-07-28 (Korf, 2004), and Exonerate version 2.2.0 (Slater & Birney, 2005). For this MAKER prediction we used as input datasets the *F. exsecta* transcriptome (ESTs) (Bioproject ID: PRJNA213662, (Dhaygude et al., 2017)), and the proteomes of all available ant species (Uniprot download on 20-04-2015). The longest protein at each genomic locus was retained, resulting in a set of 23,517 gene models. Because samples may have different sets of transcripts, owing to different biological conditions or developmental stages (Dhaygude et al., 2017), we additionally made a separate transcript-spliced assembly using RNA sequences generated from separate libraries for different life stages (Dhaygude et al., 2017), using the Tophat version 2.1.0 (Trapnell, Pachter & Salzberg, 2009), and Cufflinks version 2.2.1 (Trapnell et al., 2010). The assemblies from the different samples were then merged using cuffmerge (Trapnell et al., 2010). We further obtained separate Augustus version 3.1.0 (Stanke & Morgenstern, 2005), and Glimmer version 3.02 (Salzberg et al., 1998) gene models with default settings (Augustus: --species=fly -- genemodel=partial, --strand=both, Glimmer: +f, +s, −g 60). The gene sets and gene models from MAKER and from other programs were then merged. Redundancy was removed by favoring for each transcript the longest prediction starting with a methionine. If several transcripts had the same length we retained the one which had the best support from the cufflinks transcript assembly. This redundancy removal resulted in a final set of 13,637 protein coding gene models (final OGS), which contained 33,121 transcripts.

### Genome Annotation

We analyzed the complete official gene sets (OGS) of *F. exsecta* to identify sequence and functional similarity by comparing with different sequence databases using blast. By using a ribosomal database, we were able to annotate both the large (LSU), and the small (SSU) subunit ribosomal RNAs. The remaining gene sequences were used for retrieving functional information from other databases (SwissProt, Pfam, PROSITE, and COG). Gene sequences were considered to be coding if they had a strong unique hit to the SwissProt protein database (Magrane & Consortium, 2011; The Uniprot Consortium, 2017), or appeared to be orthologs of known predicted protein-coding genes from ant species based on TrEMBL (Translation of EMBL nucleotide sequence database). We also assigned putative metabolic pathways, functional classes, enzyme classes, GeneOntology terms, and locus names with the AutoFact tool (Koski et al., 2005). To further improve annotation, and for assigning biological function (e.g. gene expression, metabolic pathways), we also did orthologous searches by comparing with other Hymenoptera sequences (Wurm et al., 2009). To quantify variation in the numbers of protein family members, we performed Pfam (version 24.0) (Bateman et al., 2004) and PROSITE profile (Sigrist et al., 2010) analyses on proteins obtained from the *F. exsecta* gene set. Our final annotation included gene sequences with retrieved protein-related names, functional domains, and expression in other organisms along with enzyme commission (EC) numbers, pathway information, Cluster of Orthologous Groups (COG), functional classes, and Gene Ontology terms.

### Orthology and evolutionary rates

Comparative genome-wide analysis of orthologous genes was performed with OrthoVenn (Wang et al., 2015) to compare the predicted *F. exsecta* protein sequences with those of four other ant species, *Camponotus floridanus*, *Lasius niger*, *Solenopsis invicta*, and *Cerapachys biroi*, all of which were downloaded from their respective public NCBI repositories. The predicted proteins of *F. exsecta* and the other four species were uploaded into the OrthoVenn web server for identification and comparison of orthologous clusters (Wang et al., 2015). Following clustering, orthAgogue was used for the identification of putative orthology and inparalogy relationships. To deduce the putative function of each ortholog, the first protein sequence from each cluster was searched against the non-redundant protein database UniProt using blastp 2.2.27. Pairwise sequence similarities among protein sequences were determined for all species with a blastp 2.2.27 (E-value cut-off of 10^−5^, and an inflation value of 1.5 for MCL). Finally, an interactive Venn diagram, summary counts, and functional summaries of clusters shared between species were visualized using OrthoVenn.

To identify genes under positive or relaxed purifying selection in *F. exsecta*, we estimated the rates of non-synonymous to synonymous changes for core orthologous genes (3,156) from five ant species (*F. exsecta*, *Camponotus floridanus*, *Lasius niger*, *Solenopsis invicta*, and *Cerapachys biroi*). For this we only included orthologous groups with one ortholog for each species (no paralogous genes were included) in the analysis. We extracted coding and protein sequences for 3,156 orthologous groups from the respective public NCBI repositories for the species included. We then aligned all protein sequences using Clustal Omega (Sievers & Higgins, 2014), and then converted them to nucleotide sequences with PAL2NAL version 14 (Yang, 1997). We then ran CODEML version 4.9e (Yang, 1997), using the branch site model with *F. exsecta* as foreground branch, and the other five ant species as background lineages. The Bayes empirical method (Yang et al. 2005) was used to estimate the posterior probabilities, which were then used to identify sites under selection. We additionally estimated pairwise dN/dS ratios for orthologous genes (5,148 genes) between *Camponotus floridanus* and *F. exsecta* in CODEML.

We also ran an orthology analysis between the proteins from three *Wolbachia* species published previously (wRi, wDac, wNo; (Klasson et al., 2009b; Ellegaard et al., 2013; Ramirez-Puebla et al., 2016)), to find similarity with the predicted protein sets of the newly assembled wFex genome. Orthologs were identified using OrthoVenn (E-value cut-off of 10^−5^ and inflation value 1.5). In addition, we analyzed the paralogous genes within the wFex genome, to help understand the increased genome size in comparison to other *Wolbachia* genomes.

### Discovery and annotation of transposable elements

We used RepeatMasker version 4.0.7 (Smit. et al., 2015), and the TransposonPSI version 08-22-2010 (Brian J. Haas, 2011) to detect repetitive elements in the genome. To retrieve and mask repetitive elements, we downloaded files from the Repbase and Dfam databases, and aligned each of them with the *F. exsecta* genome sequences as query sequences. Positive alignments were regarded as repetitive regions and extracted for further analysis. To identify genome sequence region homology to proteins encoded by different families of transposable elements, we used the TransposonPSI analysis tool. This tool uses PSI-blast, with a collection of retro transposon ORF homology profiles to identify statistically significant alignments.

### *Wolbachia* phylogeny

We analysed the phylogeny of *Wolbachia* in MrBayes v3.2.6 x64 (Ronquist & Huelsenbeck, 2003), using a concatenated sequences of 35 genes. For this analysis, each gene was considered as a different partition, and the most fitting nucleotide substitution model was chosen for each gene, using the bayesian information criterion (BIC) in the program jMODELTEST (Posada, 2008). The partitioned dataset was run for 200,000 generations, sampling at every 100th generation with each partition unlinked for the substitution parameters. Convergence of the runs was confirmed by checking that the potential scale reduction factor was ~1.0 for all model parameters, and by ensuring that an average split frequency of standard deviations < 0.01 was reached (Ronquist & Huelsenbeck, 2003). The first 25% of the trees were discarded as burn-in, and the remaining trees were used to create a 50% majority-rule consensus tree, and to estimate the posterior probabilities. To check for consistency of the phylogeny, Markov chain Monte Carlo (MCMC) runs were repeated to get a similar 50% majority-rule consensus tree with high posterior probabilities. The phylogenetic tree generated was visualized using Figtree v1.4.2 (Rambaut, 2012).

## Results & Discussion

### Assembly of the *Formica exsecta* genome

We created Illumina sequencing libraries from DNA extracted from testes of males of a *F. exsecta* colony to obtain >99 gigabases of Illumina sequence data. The final *F. execta* genome resulting from assembly of this data was 277.7 megabases (Mb) long, encompassing 14,617 scaffolds (Figure 1) with a N50 scaffold length of 997.7 kb (Table 2). The number of scaffolds is higher than the number of chromosomes reported for *F. exsecta* (n=26; Agosti & Hauschteck-Jungen, 1987; Rosengren, Rosengren & Söderlund, 2009). Similarly, the *F. exsecta* genome assembly is somewhat shorter than genome size estimates obtained by flow cytometry for species in the subfamily Formicinae (range: 296-385 Mb; Tsutsui et al., 2008). These discrepancies are unsurprising given the difficulty of assembling highly repetitive gene content from short sequencing reads (Henson, Tischler & Ning, 2012). In line with this, the genome assembly length metrics are similar to those of the 23 ant genomes that have been published. The raw data, gapped scaffolds, and annotations underpinning this assembly are deposited into public databases under BioProject PRJNA393850 (accession NPMM00000000).

**Figure 1.**
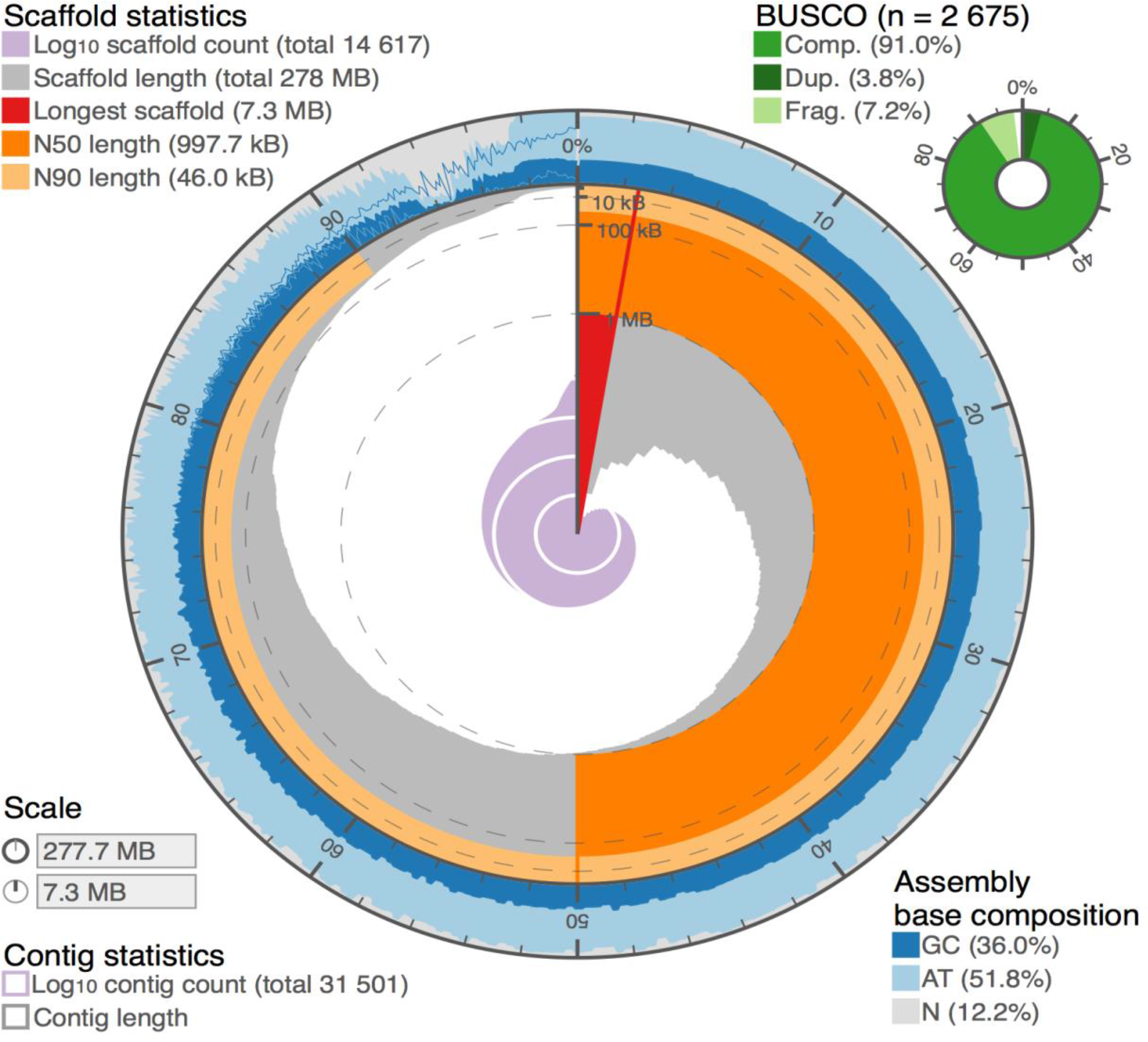
*De novo* genome assembly of *F. exsecta* genome, summarized by the following metrics: a) Overall assembly length, b) Number of scaffolds/contigs, c) Length of the longest scaffold/contig, d) Scaffold/contig N50 and N90, e) Percentage GCs and percentage Ns, f) BUSCO completeness, g) Scaffold/contig length/count distribution.

### Quantitative assessment of genome assembly

Based on scaffold N50 and N75 statistics, contig size, and GC content, the *F. exsecta* genome assembly is comparable in quality and completeness to other sequenced ant genomes (Supplementary Table S1). All the 248 CEGMA eukaryotic core genes were found, and 241 of these genes were complete in length. Similarly, 98.5% of 1634 BUSCO Insecta genes were complete in the genome (Table 3). These results held with other BUSCO analysis levels including Eukaryota, Arthropoda, and Hymenoptera, with low duplication levels (2.2% to 5.3%), and few missing genes (0.6% to 1.27%; all details in Table 3). Such discrepancies can be due to technical artifacts such as sequencing biases or assembly difficulties, as well as to true differences between our *F. exsecta* sample and the BUSCO and CEGMA datasets. To further evaluate genome completeness, we compared the independently generated *F. exsecta* transcriptome (Dhaygude et al., 2017) to the genome reported here. More than 98.75 % of the 10.999 assembled ESTs mapped unambiguously to the genome (blastn E < 10^−50^). Together, these analyses show that the genome assembly has high completeness.

### Gene Content in the *Formica exsecta* genome

We identified 13,637 protein coding genes by combining *ab initio*, EST-based, and sequence similarity based gene predictions methods. The GC content was higher in exons (41.6%) than in introns (30.6%), a pattern similar to that reported in the honey bee, *Apis mellifera*, and the fire ant, *Solenopsis invicta* (Weinstock et al., 2006; Wurm et al., 2011). Despite this, as in other ant genomes (Schrader et al., 2014; Boomsma et al., 2017), overall GC content in genes (35.1%) was similar to the rest of the genome (36.0%).

We used blast and orthology analyses to characterize *F. exsecta* genes. The vast majority (88%; 12,050) of these had the highest blastp similarity to genes in other ants. A further 0.4% had the highest similarity to Apidae, and 0.6% to Braconidae, Amniota, and *Wolbachia* (the latter probably due to HGT; see below and Figure 2). The remaining 3.09% belong to other taxa not included in Figure 2 because they had fewer than 20 hits. The remaining genes (7.91%, n= 1,080) lacked clear sequence similarity [cutoff for blastx E < 10^−3^] to known protein sequences or protein domains. Some of these may represent erroneous gene predictions (Drăgan et al., 2016), however 994 of them are ≥1000 bp and include an open reading frame >300 amino acids long, which is unlikely to occur by chance. Importantly, although only a single pooled transcriptome library, prepared from different developmental life stage samples, was available for *F. exsecta*, 235 of the genes are expressed (FPKM ≥ 1; Dhaygude et al., 2017). It is thus likely that a high proportion of the 1,080 genes are taxonomically restricted genes unique to the *F. exsecta* lineage.

**Figure 2.**
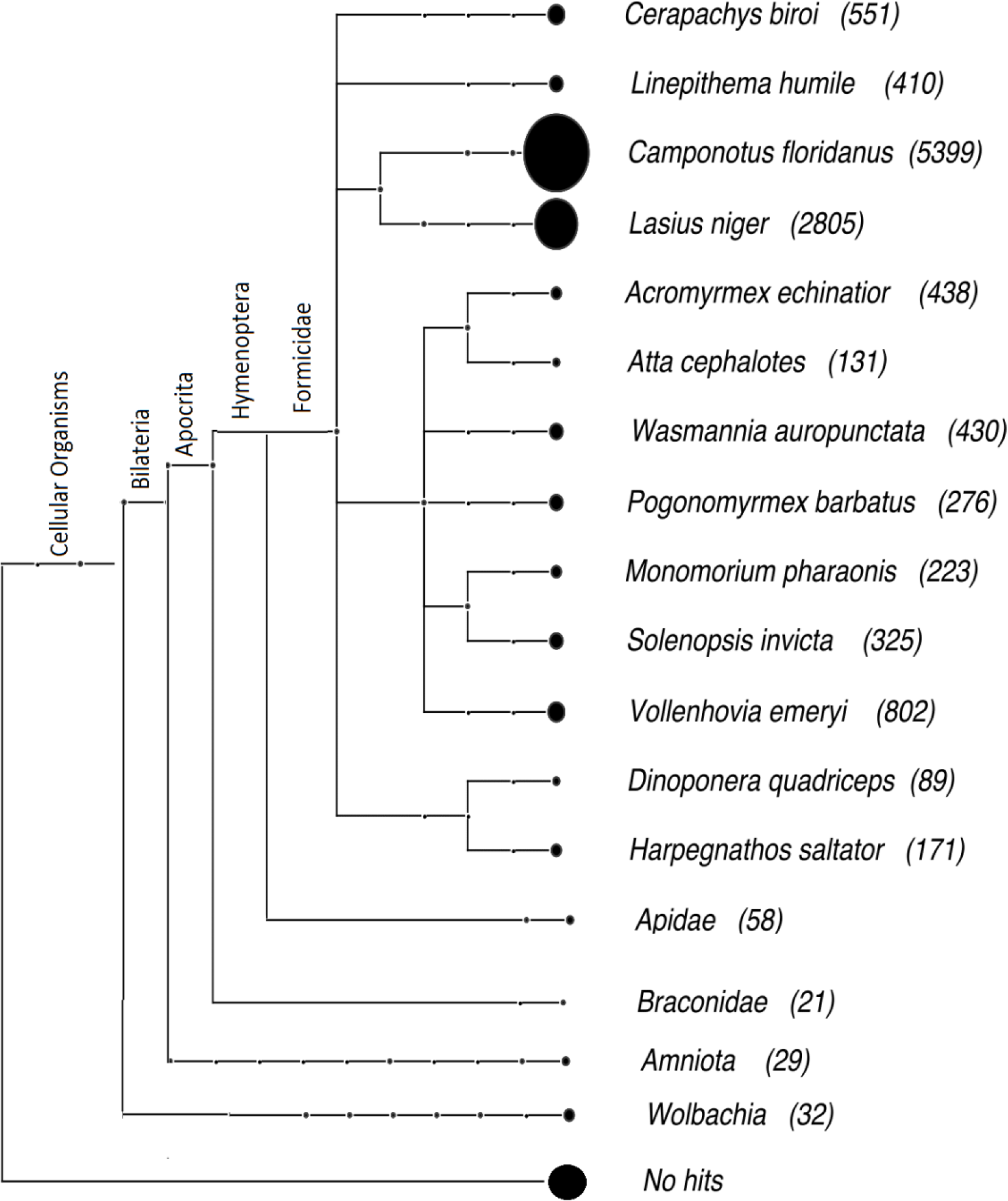
Taxonomic distribution of the best blastp hits of *F. exsecta* proteins to the non-redundant (nr) protein database (E < 10^−5^).

The total genes of *F. exsecta* (n=13,637) were grouped into 7,727 orthologous clusters (Figure 3). Comparative analysis of the *F. exsecta* genes with the closely related species *C. floridanus* and *L. niger*, and the more distantly related *S. invicta* and *C. biroi* revealed, that 4,685 orthologous clusters out of 7,727 are shared between all five species. In addition, we found 102 gene clusters that were exclusive to three Formicinae genomes (*F. exsecta*, *C. floridanus* and *L. niger;* Supplementary Table S2). Such genes are important candidates that could be involved in the evolution of this subfamily. Many of the genes in these clusters had no detectable relation to existing genes outside the Formicinae; those that did included GO annotations such as glycerate kinase, transferase activity, deoxyribonucleoside diphosphate metabolic process.

**Figure 3.**
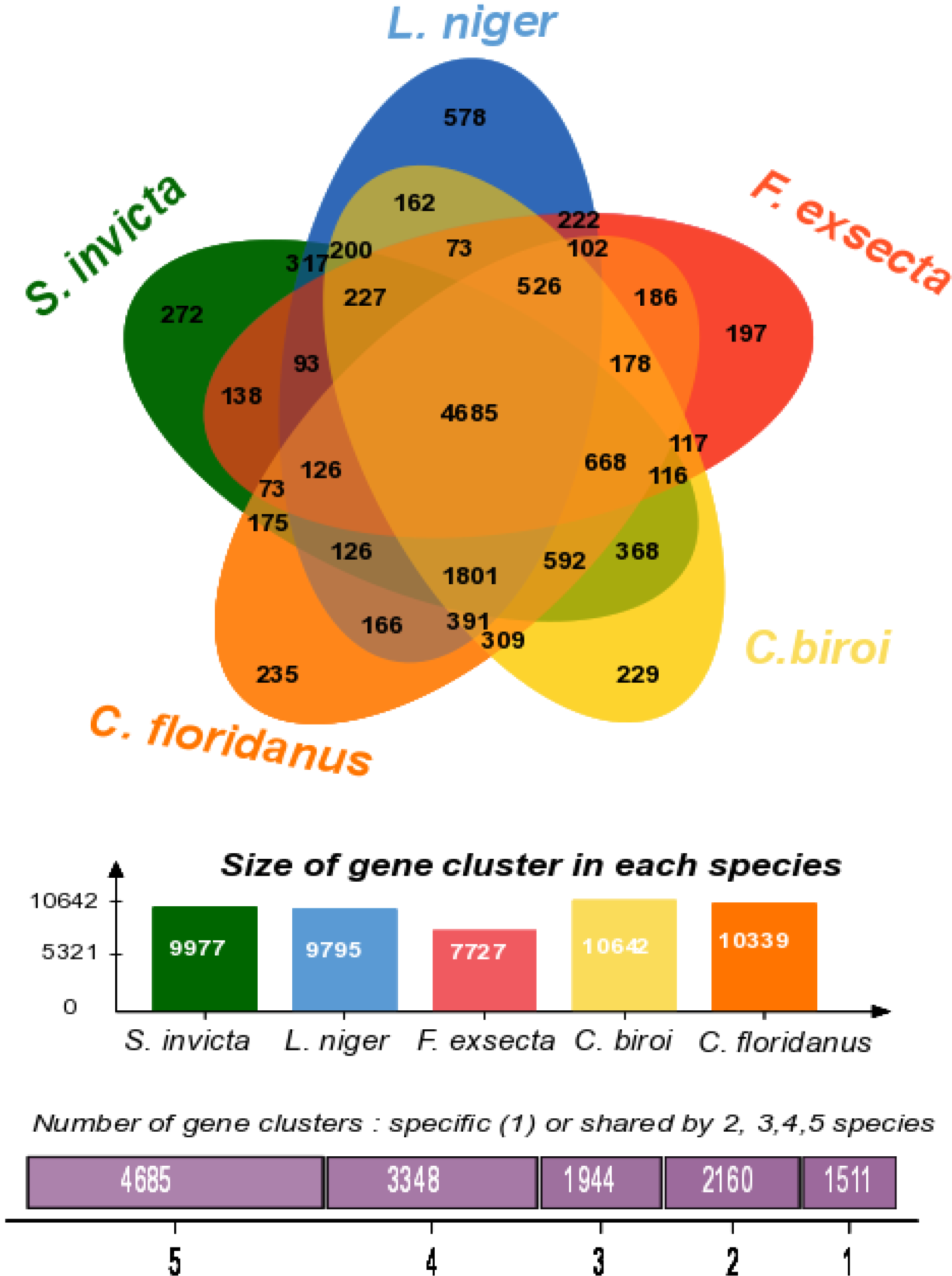
Venn diagram showing the distribution of gene families (orthologous clusters) among five ant species including three closely related members of the subfamily Formicinae (*Formica exsecta, Camponotus floridanus, Lasius niger*), and two distinctly related ants (*Solenopsis invicta* and *Cerapachys biroi)*.

Interestingly, 633 of the *F. exsecta*-specific genes could be grouped into 197 ortholog clusters of 2 or more genes (Supplementary Table S3), suggesting not only newly evolved genes, but also potential gene duplication and subfunctionalisation. Previous comparative genome studies have indicated that 10-20% of genes lack recognizable homologs in other species in every taxonomic group so far studied (Wilson et al., 2007; Khalturin et al., 2009; Johnson & Tsutsui, 2011; Tautz & Domazet-Lošo, 2011). Our lower percentage of orphan genes could be due to our hierarchical approach to annotation, the wide range of databases used, and the large amounts of ant genomic data generated over the past years (Favreau et al., 2018).

### Genes with signatures of evolution under positive selection

We performed analyses to detect genes with signatures of positive selection in *F. exsecta.* First, selection analysis (dN/dS ratio estimations) on 3,157 single-copy genes shared between the five core ant species (without paralogous genes), revealed that 500 genes have signatures of positive selection in the lineage leading to *F. exsecta*. These include genes involved in fatty acid metabolism, lipid catabolism, and chitin metabolism (Supplementary Table S4). Interestingly, previous studies on ants, bees, and flies also provide evidence for positive selection on genes in similar functional categories as in our study (Roux et al., 2014). For example, genes involved in biological functions such as carbohydrate metabolic processes, lipid metabolic processes, cytoskeleton organization, cell surface receptor signaling pathways, and RNA processing were overrepresented in the enrichment analysis, and such genes were also previously reported as positively selected genes in ants, bees, and flies (Viljakainen et al., 2009; Roux et al., 2014).

To perform a similar analysis on a larger number of genes, we used a second approach based on pairwise comparisons between *F. exsecta* and *C. floridanus*. Out of 5,148 one-to-one orthologs, 29 showed dN/dS > 1 (P < 0.005; Supplementary Table S5). Although some of these putative genes could be artefactual or non-coding, they all include an open reading frame of > 100 amino acids. Five (17%) out of 29 genes are likely linked to transposon activity as they are transposase-like or have EpsG domains. Among the other genes, only a few are annotated: the Icarapin-like protein is a venom gene, and such genes have been shown to be under positive selection in wasps (Werren et al., 2010). Perhaps more surprisingly we found high dN/dS for the Homeobox protein gene orthopedia which is involved in early embryonic development (Mackenzie et al., 1991).

### Repetitive elements

Repetitive elements comprised 15.88% (44.10 Mb) of the *F. exsecta* assembly. This proportion is similar to that found in other ants (16.5-31.5% (Schrader et al., 2014). This is probably an underestimate because (i) genomic regions that cannot be assembled are enriched with such repeats, (ii) multiple copies of a repetitive element are often collapsed into a single copy during genome assembly, and (iii) only a portion of repetitive elements in *F. exsecta* will have similarity to sequences in standard repeat databases. Overall, 3.18% (8.8 Mb) of the assembly was composed of simple repeats, whereas 12.73% (35.34 Mb) comprised interspersed repeats, most of which (53.73%) could not be classified. Among those that could be classified, 10,542 retro element fragments represented 2.74% of the genome, and 53,438 DNA transposons represented 4.23% of the genome. The *F. exsecta* genome contains copies of the piggyBac transposon (23 in total, and 7 within intact ORFs). Higher numbers (234) of piggyBac transposons have been found in *C. floridanus*, yet only 6 of these were found within ORFs (Bonasio et al., 2010).

### The *Wolbachia* endosymbiont genome of *Formica exsecta*

The assembly of the *Wolbachia* endosymbiont, wFex, was 3.09 Mb long, encompassing 69 scaffolds with a N50 scaffold length of 104,167 nt, and a GC content of 35.13% (Table 2; GenBank: RCIU00000000, Bioproject: PRJNA436771). This assembly of wFex shows extensive nucleotide similarity with the complete genome of the *Wolbachia* endosymbiont of *Drosophila simulans*, wNo (GenBank ID: NC_021084), and covers approximately 84% of its length (Supplementary Figure S2). We determined that 549 genes are present as a single copy in the *Wolbachia* genomes most closely related to wFex ((Lindsey et al., 2016) see below); 537 (99.6%) out of these 539 core genes are present in the wFex genome, suggesting high completeness.

However, the wFex genome is considerably larger (3.09 Mb) than the *Wolbachia* genomes reported previously (range: 0.95 to 1.66 Mb; Sun et al., 2001), and includes a greater number of open reading frames (1,796 ORFs) than other published *Wolbachia* genomes [range: 644 to 1,275 genes]. *Formica exsecta* is known to harbor more than one *Wolbachia* strain (Reuter & Keller, 2003), thus these patterns could be due to the presence of multiple endosymbiont strains. Two additional lines of evidence support this idea. First, 212 genes (11.80 %), that are present as single-copy genes in the wMel, wRi and wDac genomes (Klasson et al., 2009b; Ellegaard et al., 2013; Ramirez-Puebla et al., 2016), are duplicated in our assembly (Supplementary Table S6). Furthermore, 92 (12%) of the 775 genes present as a single copy in wFex, included genetic variation within our sample, including in the cytochrome oxidase subunit I; no such variation is normally expected. Despite extensive attempts, we were unable to disentangle the two or more *Wolbachia* strains - this is likely because differences in synteny between the strains cannot be resolved using short-read sequence data. Similar assembly artifacts, due to multiple *Wolbachia* strains, have also been reported by other studies (Ramírez-Puebla et al., 2016).

To determine how wFex is related to other *Wolbachia*, we used Bayesian phylogenetic analysis based on 35 conserved genes (Supplementary Table S7) from the 25 available *Wolbachia* genomes from the NCBI database. The analysis revealed three distinct monophyletic clades, all with posterior probabilities >0.9. Each of these clades represent one super group of *Wolbachia* (Figure 4). Of these three supergroups, two have been found only in arthropods (super groups A and B), and the third super group is found only in filarial nematodes (super group C; Werren, Baldo & Clark, 2008). In the phylogenetic analysis, wFex clustered with the *Wolbachia* strains within super group A, and most closely matched the strain that infects the scale insect, *Dactylopius coccus*, (wDacA). This is consistent with earlier studies on *Wolbachia* in ants, which also found supergroup A in the majority of the infected ants (Werren & Windsor, 2000). Given that wFex affiliates with the supergroup A in our phylogenetic analysis, we investigated the extent to which its gene content aligned with that of other *Wolbachia* genomes in the same supergroup. We found that 525 genes were shared across all strains in this supergroup, including wFex (Figure 5). About 20% of these genes had no match to known proteins, whereas the remaining genes matched a wide range of predicted functions (Ellegaard et al., 2013; Lindsey et al., 2016). We also found strain-specific genes (wFex - 50 genes, wMel - 4 genes, wRi - 3 genes, wDac - 9 genes). The wFex-specific genes included inferred annotations including Ankyrin repeat protein, ATP synthase, and chromosome partition protein (Supplementary Table S8). These strain-specific genes can provide an interesting snapshot of the evolutionary dynamics of a species. For example, ankyrin repeat proteins are involved in numerous functional processes, and have been suggested to play an important role in host-symbiont interactions (Li, Mahajan & Tsai, 2006). Comparative analyses suggest that they may be involved in host communication and reproductive phenotypes (Voronin & Kiseleva, 2008).

**Figure 4:**
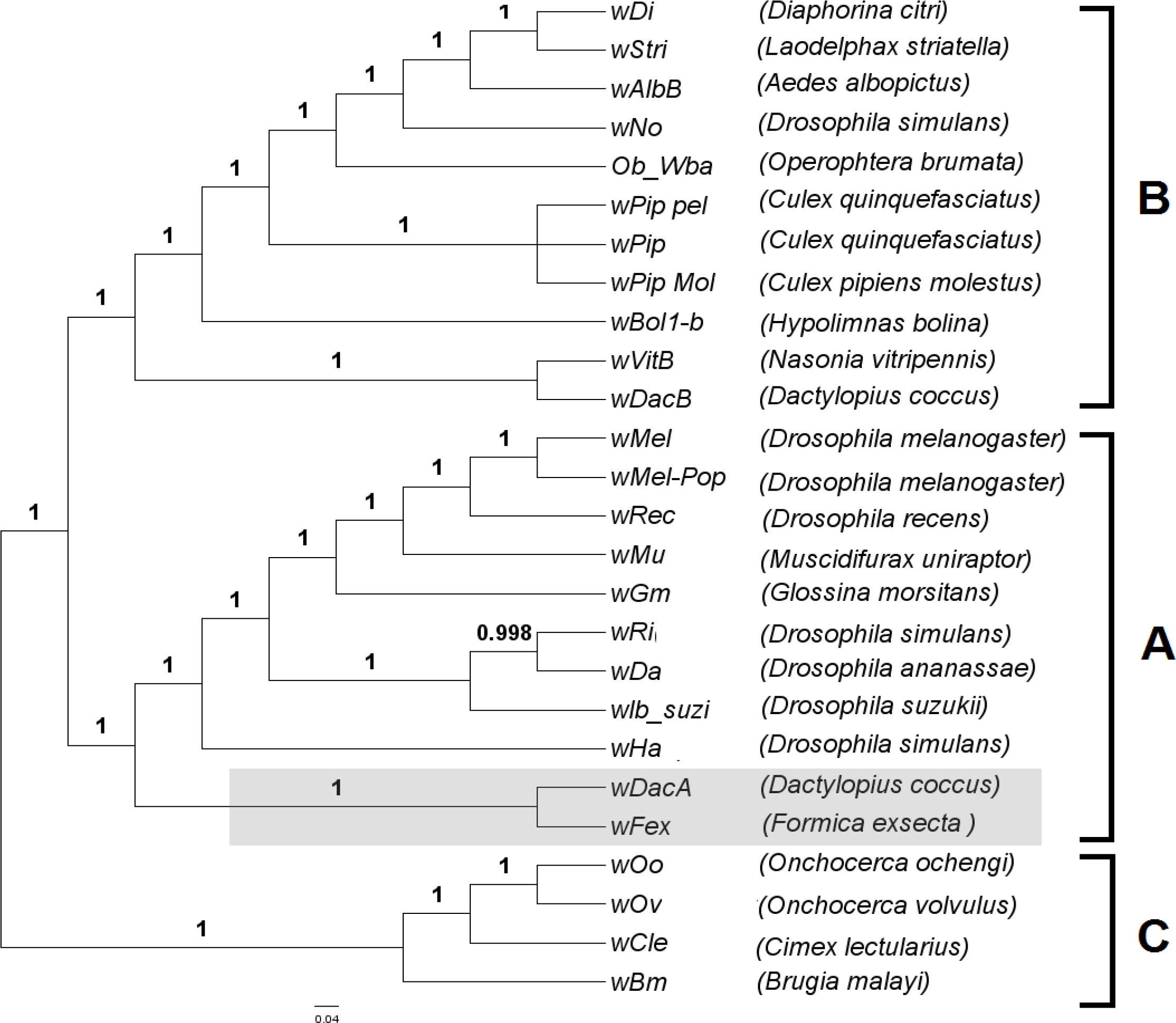
Phylogeny of the *Wolbachia* supergroups A, B, and C strains with the newly assembled wFex genome. The phylogenetic reconstructions are based on individual analyses of 35 core genes of 25 *Wolbachia* strains. The support values on the branch labels indicate Bayesian posterior probabilities. The letters A-C indicate the separate supergroups.

**Figure 5.**
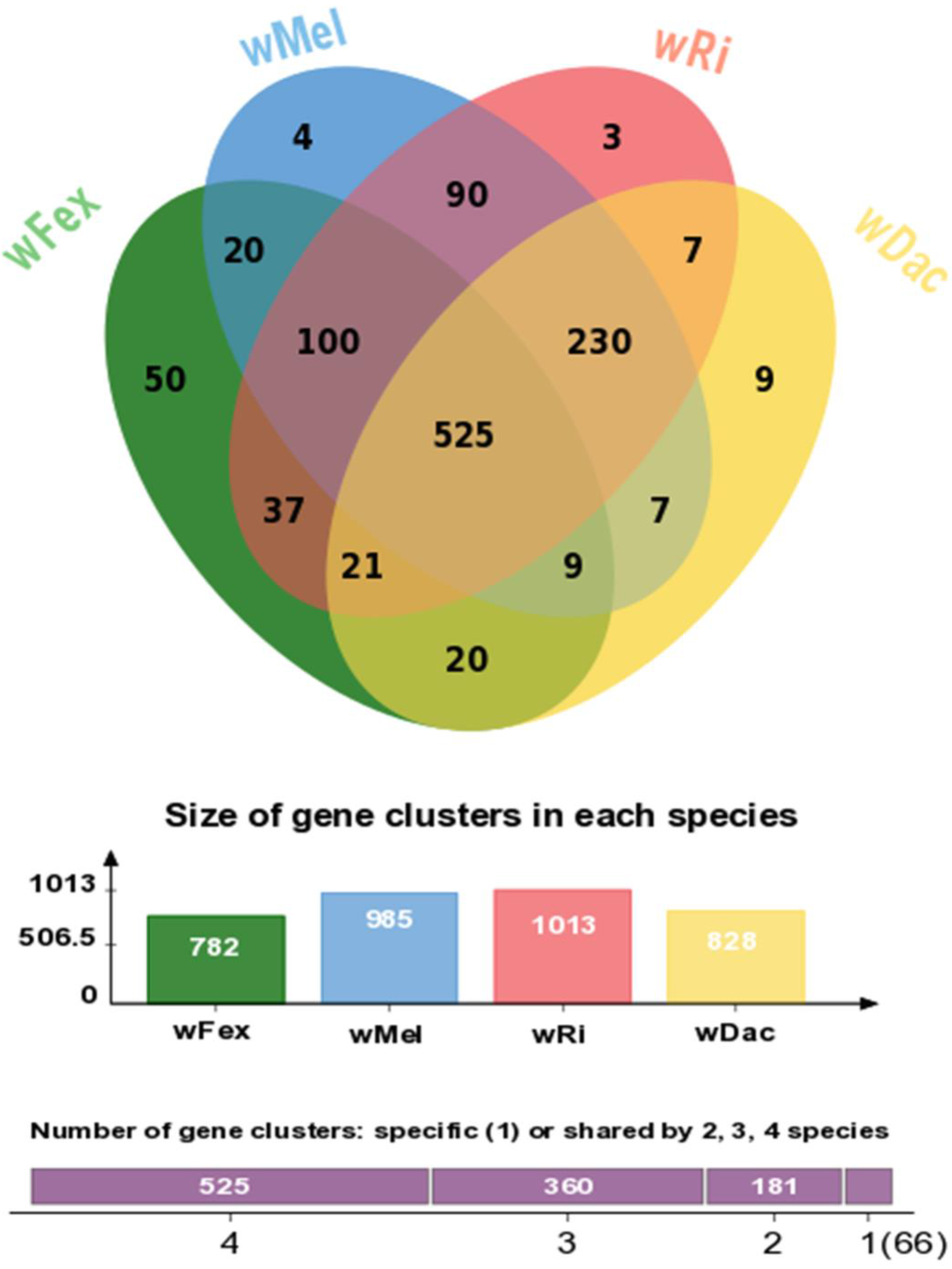
Venn diagram displaying the overlap in orthologous genes among four *Wolbachia* species including the newly assembled wFex strain and the wDac, wRi, wMel strains reported previously.

To explore differences in gene content between CI-inducing and mutualist strains of *Wolbachia*, homologous genes in six CI-inducing strains, and three mutualist strains were aligned and compared (Lindsey et al., 2016). The mutualist *Wolbachia* strains (range: 644-805 genes) had fewer genes than the CI-inducing ones (range: 911-1,275 genes). The CI-inducing strains shared 84 genes not found in the mutualist strains. We found 80 (95.23%) of these 84 genes in wFex (Supplementary Figure S3), suggesting that wFex may be CI-inducing.

### Horizontal gene transfers, and functional novelty

Intracellular symbionts can contribute new genes or fragments of genes to the host genome via horizontal gene transfer (Keeling & Palmer, 2008; Werren, Baldo & Clark, 2008; Dunning Hotopp, 2011). We found evidence for ancestral horizontal transfer from *Wolbachia* to the host *F. exsecta* in five scaffolds (scaffold83, scaffold233, scaffold574, scaffold707, scaffold741). The four largest transfers are 13 to 47 kb long, and include 83 putative functional protein coding genes, whereas the fifth and smallest insertion (475 bp) lacks protein coding genes other than a degenerate *Wolbachia* transposase. This transposase is present in 7 out of 29 published *Wolbachia* genomes. Our analysis shows that similar transfer events of this homologous fragment apparently also have occurred from *Wolbachia* to the genomes of the ants *Vollenhovia emeryi* (gene: LOC105557741), and *Cardiocondyla obscurior* (scaffolds scf7180001101632 and scf7180001108526), as well as the microfilarial nematode *Brugia pahangi*, the Arizona spittle bug *Clastoptera arizona*, and the parasitoid wasp *Diachasma alloeum*.

One-third of invertebrate genomes are thought to contain recent *Wolbachia* gene insertions, ranging in size from short segments (<600 bp), to nearly the entire genome (Hotopp et al., 2007; Werren, Baldo & Clark, 2008). Most of these transferred fragments contained transposable elements, as well as some other functional genes from the *Wolbachia* genome. The HGT events from *Wolbachia* to *F. exsecta* are located in or near regions with transposases. Our blast results suggest that four of the insert regions had *Wolbachia* transposases, whereas one insert region has a transposase of ant origin. Whether the presence of such transposases close to HGT sites facilitates insertions is unknown. Interestingly, the putative functional protein coding genes of *Wolbachia* inserted in the *F. exsecta* genome are similar to the genes reported in similar HGTs events in other insect genomes (eg: ABC transporter, Ankyrin repeat containing protein (Table 4) (Brelsfoard et al., 2014; International Glossina Genome Initiative, 2014). This could indicate that some HGT events are either more likely to occur or to be retained for reasons that could be neutral or adaptive to the host or to the endosymbiont. The transcriptome of *F. exsecta* shows that at least 6 out of the 83 genes from the *Wolbachia* HGT regions are transcribed but with a low FPKM values (range 0.04 to 1.6). These low level transcription trait often observed in bacteria-eukaryote HGTs (Hotopp et al., 2007; Nikoh et al., 2008; Dunning Hotopp, 2011).

**Table 4:** HGT inserts from *Wolbachia* present in the genome of *F. exsecta* with details of length and position in the *F. exsecta* genome. The presence of similar insert regions in other eukaryote genomes is also shown.

| *Wolbachia* gene name | HGT region in *F. exsecta* | Length HGT (bp) | Transposon region near HGT | Transposon Name | Observed in other species | Other Host Species name with position of similar insertion |
| --- | --- | --- | --- | --- | --- | --- |
| Transposase | scaffold83: 2271642-2272117 | 475 | scaffold83:2271642-2272117 | transposase | Complete | *Vollenhovia emeryi* (LOC105557741), *Cardiocondyla obscurior* (genes: scf7180001101632 and scf7180001108526), *Diachasma alloeum* (LOC107035412), *Brugia pahangi* (BPAG_contig0001587), |
| ABC transporter ATP-binding protein, porphobilinogen deaminase, D-alanine--D-alanine ligase, DNA processing protein DprA , triose-phosphate isomerase | scaffold233: 1712452-1725498 | 13046 | scaffold233:1714122-1714241 | transposase | Partial (few gene region) | *Vollenhovia emeryi*(NW_011967015.1,NW_011967060.1), *Wasmannia auropunctata* (scf7180000683207,scf7180000730160), *Rhagoletis zephyria*(NW_016158779.1), *Planococcus citri*(KF021963.1) , *Ctenocephalides felis* (KC177865.1) |
| DNA repair protein RadC,transposase,DNA ligase,ABC transporter permease,ATP-dependent protease La | scaffold574: 102007-116197 | 14190 | scaffold574:105963-106483 | transposase | Partial (few gene region) | *Vollenhovia emeryi* (LOC105557101,NW_011966940.1,NW_011966751.1 ), *Monomorium pharaonis* (scf7180001140281), *Rhagoletis zephyria* (LOC108377626), *Parasteatoda tepidariorum*(LOC107444616, LOC107450900) |
| probable carboxypeptidase,type IV secretion system,conjugal transfer protein TrbL,lysyl-tRNA synthetase,UDP-N-acetylmuramoylalanine-D-glutamate ligase | scaffold707: 1-38814 | 38813 | scaffold707:35826-36154 | Mariner Mos1 transposase (Ant origin) | Partial (few gene region) | *Vollenhovia emeryi* (NW_011966954.1,NW_011966496 ), *Wasmannia auropunctata* (scf7180000735528), *Brugia pahangi* (BPAG_contig0000608, BPAG_scaffold0000225) |
| DNA methylase, Ankyrin repeat domain protein, regulatory protein RepA,site-specific recombinase, cytochrome b-like | scaffold741: 1-47265 | 47264 | scaffold741:54020-54482, scaffold741:52587-52910 | IS110 family transposase, Integrase | Partial (few gene region) | *Vollenhovia emeryi* (LOC105557561,NW_011966954.1,NW_011967060.1,NW_011967015.1 ), *Wasmannia auropunctata* (LOC105460331 , scf7180000733651), *Drosophila ananassae* (WD_0580 gene) |

## Conclusions

Here we present the first draft genome of the ant *F. exsecta*, and its *Wolbachia* endosymbiont. This is the first report of a *Wolbachia* genome from ants, and provides insights into its phylogenetic position. We further identified multiple HGT events from *Wolbachia* to *F. exsecta*. Some of these have also occurred in parallel in several other insect genomes, highlighting the extent of HGTs in eukaryotes. We expect that the *F. exsecta* genome will be a valuable resource in understanding the molecular basis of the evolution of social organization in ants: Recent genomic comparisons between *Formica selysi* and *S. invicta* have shown convergent evolution of a social chromosome, that underpins social organisation in these ants (Purcell et al., 2014). Additional comparison of these genomic regions with *F. exsecta* could provide valuable insights on the evolution of genomic architectures underlying social organization.

## Acknowledgements

The authors thank Kalevi Trontti, Jenni Paviala and Minttu Ahjos for help with the laboratory work, Pekka Pamilo, Jonna Kulmuni for useful comments on an earlier draft of the manuscript. This work was funded by the Academy of Finland (Centre of Excellence in Biological Interactions, grants no. 252411 and 284666 to L. Sundström), the University of Helsinki (to L. Sundström), the Biotechnology and Biological Sciences Research Council (grant no. BB/K004204/1 to Yannick Wurm) and the Natural Environment Research Council (grant NE/L00626X/1 to Yannick Wurm).

## Data Accessibility

The raw Illumina sequences of paired-end and mate-pair libraries are deposited on the National Center for Biotechnology Information (NCBI) under the bio-project number PRJNA393850, with the accession numbers SAMN07344805-SAMN07344811. The assembled genome sequence of *F. exsecta* is deposited on Genbank with the accession number NPMM00000000. Similarly, the draft genome assembly of wFex is deposited under the project number PRJNA436771.

## Supplementary Tables

S1: Comparison of assembly statistics of the *F. exsecta* genome and 13 other published ant genomes.

S2: List of genes specific to the Formicinae as identified by OrthoVenn.

S3: List of species-specific genes in *F. exsecta*, as identified by OrthoVenn.

S4: List of *F. exsecta* genes under positive or relaxed purifying selection (dN/dS ratios > 1) in comparison to five other ant species (*Camponotus floridanus, Lasius niger, Solenopsis invicta* and *Cerapachys biroi*)

S5: List of *F. exsecta* genes showing dN/dS ratios > 1 in pairwise comparison to *Camponotus floridanus.*

S6: List of genes with paralogs in the wFex genome, which are present as single copies in the wMel, wRi, wDac genomes.

S7: List of conserved *Wolbachia* genes used for phylogenetic analysis.

S8: List of species-specific genes in wFEX genome, as identified by OrthoVenn.

## Supplementary Figures

S1. TAGC plot of *F. exsecta*, and its *Wolbachia* endosymbiont. The TAGC plots were taxonomically annotated, and the contigs with best similarity to Arthropoda and Proteobacteria are highlighted in color.

S2. Visualization of genome coverage of wFex against the *Wolbachia* endosymbiont of *Drosophila simulans* (wNo) genome, using the alignment software Mummer.

S3. Venn diagram displaying the overlap in orthologous genes across CI-inducing and mutualist *Wolbachia* species.

